# Spatial scaling of pollen-based alpha and beta diversity within forest and open landscapes of Central Europe

**DOI:** 10.1101/2020.08.18.255737

**Authors:** Vojtěch Abraham, Jan Roleček, Ondřej Vild, Eva Jamrichová, Zuzana Plesková, Barbora Werchan, Helena Svitavská Svobodová, Petr Kuneš

## Abstract

Pollen is an abundant fossil and the most common proxy for plant diversity during the Holocene. Based on datasets in open, forest, and mixed habitats, we used the spatial distribution of floristic diversity to estimate the source area of pollen diversity and identify factors influencing the significance of this relationship.

Our study areas are Bohemian-Moravian Highlands and White Carpathians (the Czech Republic and the Slovak Republic).

Sampling 60 sites in forest and open habitats in two study regions with contrasting floristic diversity, we calculated taxonomic richness (alpha diversity) and total spatial variance (beta diversity) for pollen and floristic data along two transects, each 1 km long. Following this, we calculated the correlation between floristic and pollen diversity. We also assessed the consistency of the relationship in different habitats. Finally, we regressed local contributions of individual sites to the beta diversity of pollen and floristic data in each of the regions.

There was a positive correlation between pollen and floristic richness in both habitats in both regions; open and mixed datasets were significant. The highest correlation (adjusted R2) mostly occurred within the first tens of metres (1.5–70) and then within the first hundreds of metres (250–550). Variances of pollen data significantly correlated with variances of floristic data between 100 and 250 m. Local contributions to beta diversity of pollen and plants significantly correlated in the forest and one of the mixed datasets.

Floristic richness at the pollen site and position of the site within the landscape structure determine the sequence of the appearing species in the increasing distance. The number of species sets the source area of pollen richness and dissimilarity of appearing species controls the source area of pollen variance. These findings, linking pollen and floristic diversity, provide an essential stepping-stone for the reconstruction of historic plant diversity.

## Introduction

The ongoing, human-induced changes to biodiversity call for a progress in our understanding of past biodiversity and its long-term dynamics. Pollen is one of the most frequently used proxies of past plant diversity and composition (Birks 2019); therefore, a deeper understanding of pollen-vegetation relationship is essential. Numerous pollen–vegetation studies focusing on species composition (Davis 1963) have led to the development of spatially explicit models for plant abundances in the past (Sugita 2007, Theuerkauf and Couwenberg 2017). The fundamental paleoecological proxy of plant diversity is pollen richness, i.e., the number of pollen types in the record, but available comparisons of current pollen and floristic richness, i.e., the number of plant species in the surrounding vegetation, resulted in a positive relationship in only a few studies (Birks 1973, Meltsov et al. 2011, Felde et al. 2015, Reitalu et al. 2019, Blaus et al. 2020).

Floristic data of these studies covered different spatial scales. The local scale, corresponding to alpha diversity in ecological studies, was captured by field surveys of the surrounding landscape, either in a defined radius from the pollen sample (250 m in Meltsov et al., 2011; 500 m in Felde et al., 2015; 100 m in Blaus et al., 2020) or in clearly delimited plots (20 × 20 m in Birks, 1973). Large-scale studies usually relied on floristic data available from databases and floras, which suffer from low spatial precision. Consequently, the resulting richness corresponds to gamma diversity rather than to alpha diversity as conceived in ecological studies (grid cells 50 × 50 km; (Reitalu et al. 2019). An empirical estimate of the relevant source area of pollen (RSAP; Sugita, 1994) is frequently used to detect the spatial scale captured by the pollen record. It usually spans from ten metres (Calcote 1995) to a few thousand metres (Abraham and Kozáková 2012) around the pollen sampling site. However, the RSAP concept is designed for frequent species in the pollen spectra (Bunting et al. 2013), while pollen richness is strongly affected by rare taxa, including i) herbs originating from the nearest surroundings (Bunting 2003), and ii) extra-regional pollen component, originating from distances greater than 100 km (Janssen 1973). The relevant spatial scale of floristic richness in reference to pollen richness is nevertheless mostly unknown.

Pollen richness is strongly influenced by an abundance of high pollen producers, such as *Pinus* or *Betula* (Odgaard 1999). These taxa tend to dominate the pollen rain and decrease the probability of detection of the rare taxa, which are often represented by a single pollen grain. Application of the correction factors (Andersen 1970) derived from pollen productivity estimates helps to equalise the representation of different taxa and may lead to a stronger positive relationship between pollen and floristic richness (Reitalu et al. 2019). Indeed, a study from an altitudinal transect in southern Norway showed that the strongest representation bias appears in the boreal forest biome, which is dominated by high pollen producers (Felde et al. 2015). In this respect, studies are needed in the regions where high pollen producers do not dominate the vegetation.

Landscape structure also significantly affects pollen diversity. Open landscapes have a larger source area of pollen than forest landscapes (Hellman et al. 2009a), though both types of landscape represent extremes in alpha diversity in temperate and boreal regions. Here, forests are generally species-poorer than open landscapes, and when sites from both types of landscape are included, pollen richness is significantly regressed to floristic richness (Meltsov et al. 2011). In general terms, landscape structure corresponds to beta diversity, i.e., the variation of species composition among sites within a study area (Whittaker 1960).

There were several attempts to estimate beta diversity from fossil pollen records using various beta diversity indices. Beta diversity can be understood as a compositional turnover across time or space. Both directional metrics can be estimated between a pair of pollen assemblages by dissimilarity coefficient as a rate-of-change (e.g. Figueroa-Rangel et al. 2010), or within the set of assemblages as gradient length in ordination space (e.g. Connor et al. 2019). Turnover represents an ecological process of species replacement; however, beta diversity can also be understood as nestedness (Baselga 2010), characterising the extent to which species assemblage is a subset of a species-rich sample (Blarquez et al., 2014).

Presumably, however, only a single study found a positive relationship between pollen and vegetation in the turnover calculated from data in a grid of 60 × 60 km (Nieto-Lugilde et al. 2015). Other calibration experiments approximated beta diversity by different measures of landscape structure (Meltsov et al. 2013, Matthias et al. 2015). It therefore follows that studies treating open and forest habitats separately may provide a deeper understanding of the relationship between pollen diversity and different components of plant diversity.

Our two study areas in the temperate zone of Central Europe are dominated by semi-open landscape with forest occupied by oak or spruce, which do not belong among high pollen producers (Kuneš et al. 2019; see Appendix 1, Table A1 and Fig. A1 for further information.) We hypothesise significant regression in our datasets when open sites and forest sites are analysed together, much the same as the pattern reported by Meltsov et al. (2011). We aimed to explore the relationship between the alpha diversity (richness) and beta diversity (variance) of the pollen samples and surrounding vegetation within datasets of sites from forest, open, or both habitats together. Furthermore, we aimed to interpret the strength of the relationship whilst considering the radius of the vegetation survey.

## Material and methods

### Study area

The Žďárské vrchy Mts. are a part of the Bohemian-Moravian Highlands: the most extensive highland area in the Czech Republic. The landscape is covered mainly with *Picea abies* plantations, with patches of low-productivity grasslands and agricultural fields concentrated around villages. The area is relatively poor in plant species, and its Holocene environmental development is assumed to have been dominated by forests (Roleček et al. 2020). Sampling sites were placed over an area of 650 km^2^ at altitudes between 569 and 760 m a.s.l.

The south-western White Carpathian Mts. are situated on the periphery of a forest-steppe region of the Pannonian Basin (Rasser et al. 2008, Chytrý 2012). The mildly undulating landscape is covered by a diverse mosaic of vegetation, including patches of broad-leaved forests dominated by *Quercus robur, Carpinus betulus* and *Fagus sylvatica*, as well as mown, semi-natural dry and mesic grasslands, fields, orchards, and vineyards. The area is considered a hotspot of fine-scale plant species richness (Wilson et al. 2012, Roleček et al. 2014) and harbours many rare species with disjunct distributional ranges (Hájková et al. 2011). It is a part of the White Carpathians Protected Landscape Area and Biosphere Reserve. Sampling sites were placed over an area of 250 km^2^ at altitudes between 205 and 685 m a.s.l.

### Fieldwork

We sampled 19 sites in the forest and 20 sites in open habitats (mesic and steppic meadows) of the White Carpathian Mountains region (hereafter WCM), and 10 sites in the forest and 11 sites in open habitats (wet meadows) of the Bohemian-Moravian Highlands (hereafter BMH; see Fig. 1). Open sites were selected in continuous non-forest habitats with the minimum distance to a mature tree at least 10 m. Forest sites were located within a continuously forested area, in a forest gap of at least 1 m^2^ to reduce the gravity component of pollen fallout without the contribution of wind dispersal (Sugita 1994). We tried to avoid overlapping of sampling sites and simultaneously kept the sampled area compact and homogeneous in terms of available vegetation types and environmental conditions.

**Fig. 1:**
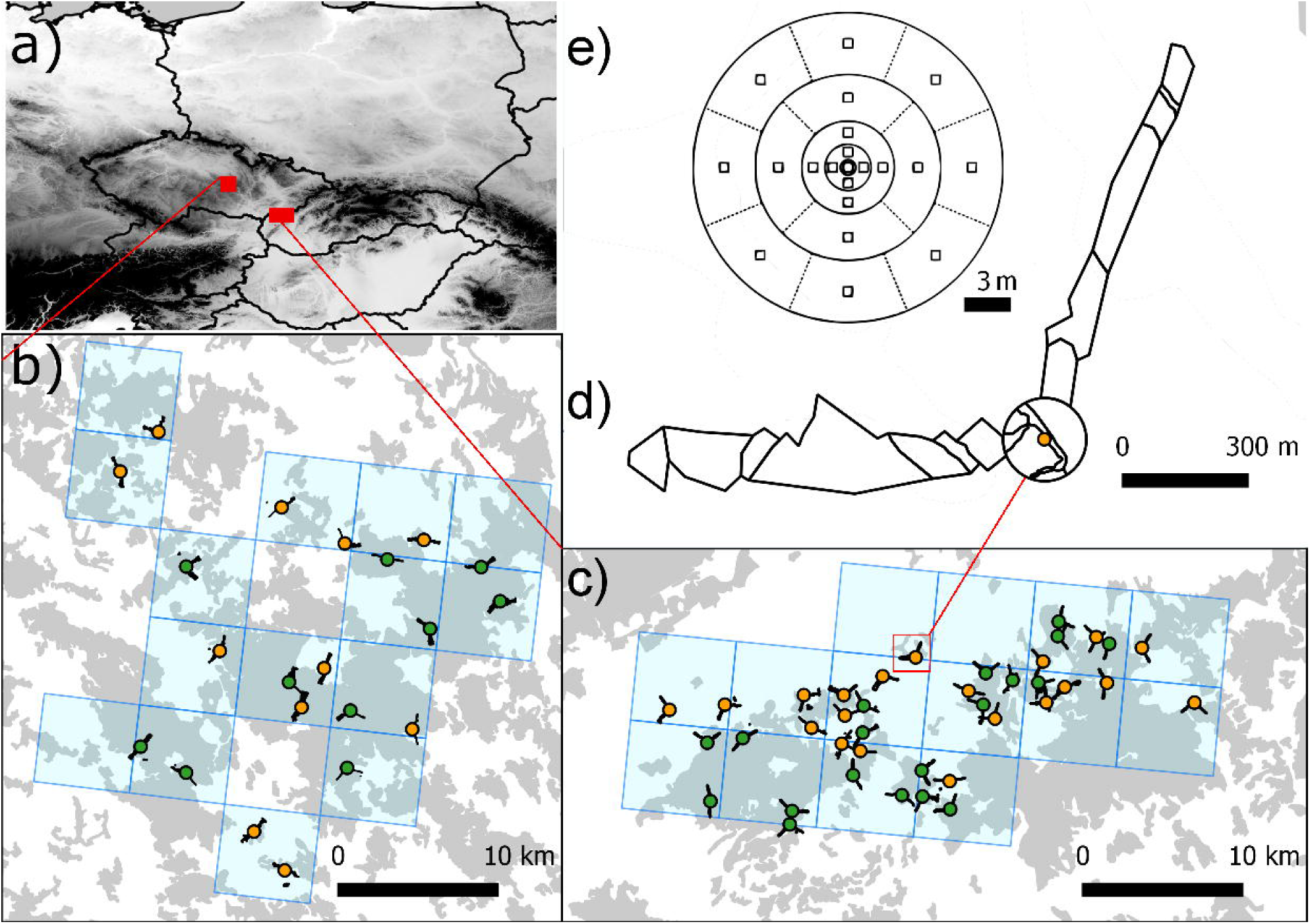
Map of the study areas showing a) position within Central Europe, b) BMH: Bohemian-Moravian Highlands, c) WCM: White Carpathians. Yellow and green circles indicate sites in meadow and forest, respectively. Blue squares show the area of the reference floristic data from the PLADIAS database (Wild et al. 2019). Grey indicates forested area. Short lines represent transects of the vegetation survey; d) circle 10–100 m and two transects of polygons recording the floristic diversity 100–1000 m, e) 21 plots within 0–10 m.

We collected pollen samples from a moss cushion of at least 50 cm^2^ in the central point of each site, while vegetation data were collected in a 1 km radius around the central point in the same year as the pollen data (Table A2). The effort of vegetation sampling was spread into three zones. Within the first 10 m, we recorded complete species lists in both regions; however, in WCM, 21 additional plots of 1 m^2^ were sampled following a modified CRACKLES protocol (Bunting et al., 2013) within the 10 m circle to evaluate finer-scale relationships. Between 10 and 100 m, main vegetation types were mapped in the field with the help of aerial photographs. The occurrence of additional species not present in the first 10 m was recorded for each of the mapped polygons. Up to 1,000 m, we recorded additional plant species, and vegetation types were mapped along two 20-metre-wide linear transects. Directions of the transects were chosen based on the aerial map to cover the highest possible habitat diversity. At the same time, the two transects had a minimum angular distance of 90°. Moreover, additional habitats not recorded in the transects were mapped and additional species recorded. Cultivated plants including ornamentals (e.g. *Thuja, Bergenia*) were also recorded.

Based on the collected data, we compiled six datasets: two “uniform datasets” with forest and open sites separated for each region, and one “mixed dataset” including both forest and open sites for each region.

### Laboratory and pollen counting

Moss polsters were prepared for pollen analysis using standard procedures (Faegri et al., 1989). Moss samples were shaken in KOH during the night, and then acetolysed for 2 min. The pollen concentrate was stored in glycerine or silicone. Pollen slides were counted under the light microscope at 400× magnification; for selected taxa at 1,000× magnification. The original pollen sum includes all pollen and spores of vascular plants following the determination key of Beug (2004).

### Numerical methods

Due to the varying pollen sum across samples (between 943 to ca. 4,000 grains), we unified the sum to 943 grains per sample by random selection without replacement and repeated this 100 times. The median number of taxa across the selections was used for further calculation. The number of pollen taxa (pollen richness) was regressed against floristic richness. Although the distances spanned from 0.5 to 1,000 m, we considered pollen and floristic richness at this scale as an alpha diversity. The concept of beta diversity in ecology is less equivocal, and there are many definitions and corresponding ways as to how to calculate beta diversity (Anderson et al. 2011). Here, we adhered to the total variance of the site-by-species community table as a measure of beta diversity (Legendre and De Cáceres 2013). Total variance represented by BDtotal value is a sum of squares in the site-by-species community table. We used the Jaccard index on presence-absence data as its measure. The relative character of BDtotal ranging from 0 to 1 allows for different numbers of sites, thus also enabling comparison between mixed and uniform datasets. Pollen BDtotal values calculated for our six datasets were regressed against six floristic BDtotals at different distances from sampling sites. Calculation of beta diversity in package adespatial (Legendre and De Cáceres 2013, Dray et al. 2020) also provided measurements of the local contribution of each site to beta diversity (hereafter also “local contribution”) and its significance. We explored the relationship between pollen and floristic counterparts at different distances from sampling sites, again by linear regression. The strength of relationships (richness, BDtotal, and local contribution) was measured by adjusted R^2^. The correlation test proved its significance at the 5% alpha level.

We used the software package R (version 3.4.3) for all statistical analyses (R Development Core Team 2017).

## Results

### Trends and ranges of richness and variance values

In both regions, we found 169 pollen types (95 in BMH and 151 in WCM) and 1,323 plant species (799 in BMH and 1,098 in WCM). Mean pollen richness per sample varied between 50 pollen types in WCM meadows, 42 pollen types in WCM forest, 38 pollen types in BMH meadows, and 31 pollen types in BMH forest (Fig. 2). Floristic richness followed the same order as pollen richness at a distance between 600 and 1,000 m: in WCM, forest species gradually increased along the whole transect, with the lowest richness between 40 and 200 m. In other datasets, however, the increase was more irregular, with more than half of the species appearing already within the first 100 m (Fig. 3).

**Fig. 2:**
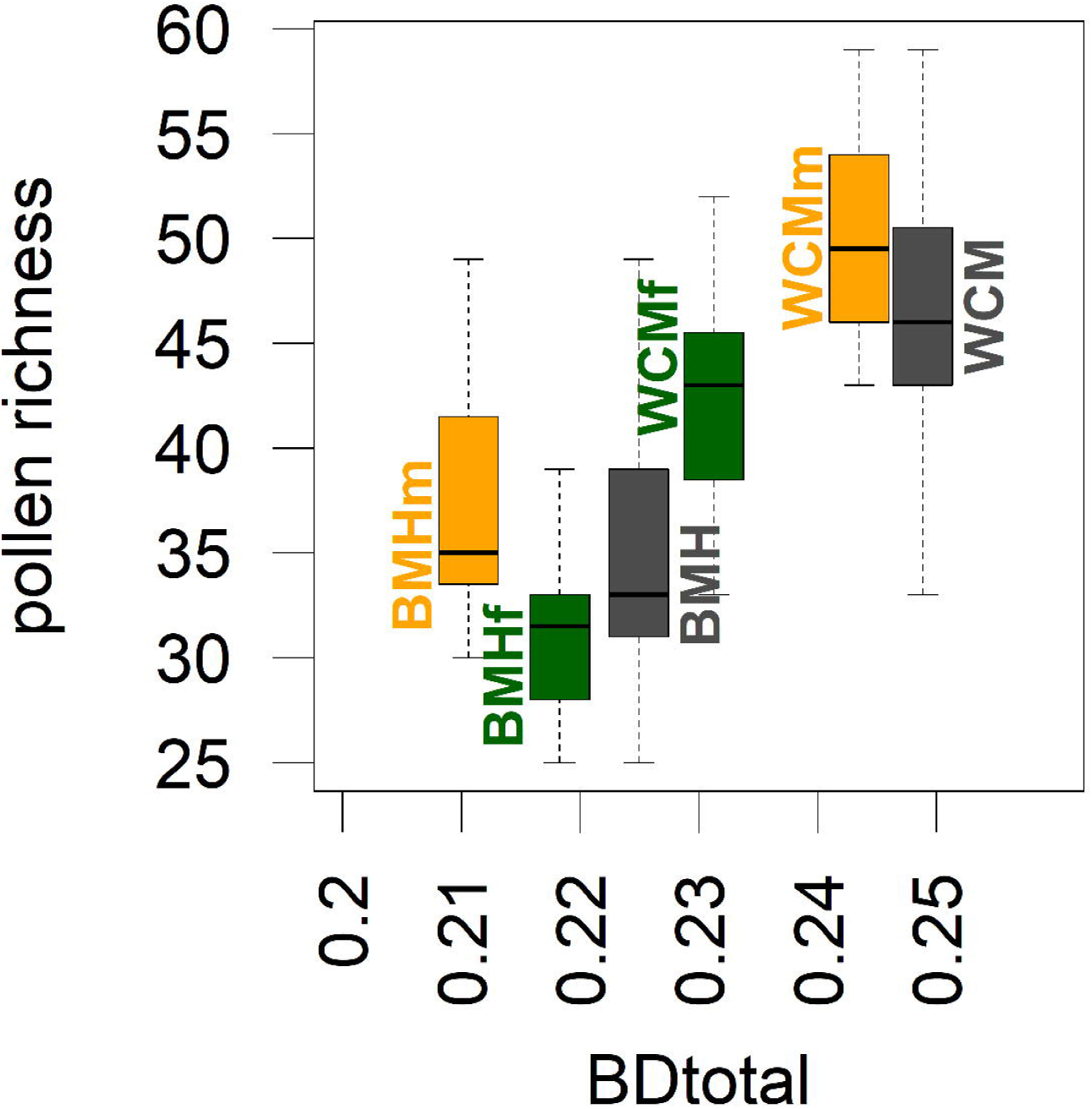
Pollen alpha diversity (pollen richness, y-axis) and beta diversity (BDtotal, x-axis) in two study regions and their different habitats. Meadows (yellow), forest (green), and both habitats together (black).

**Fig. 3:**
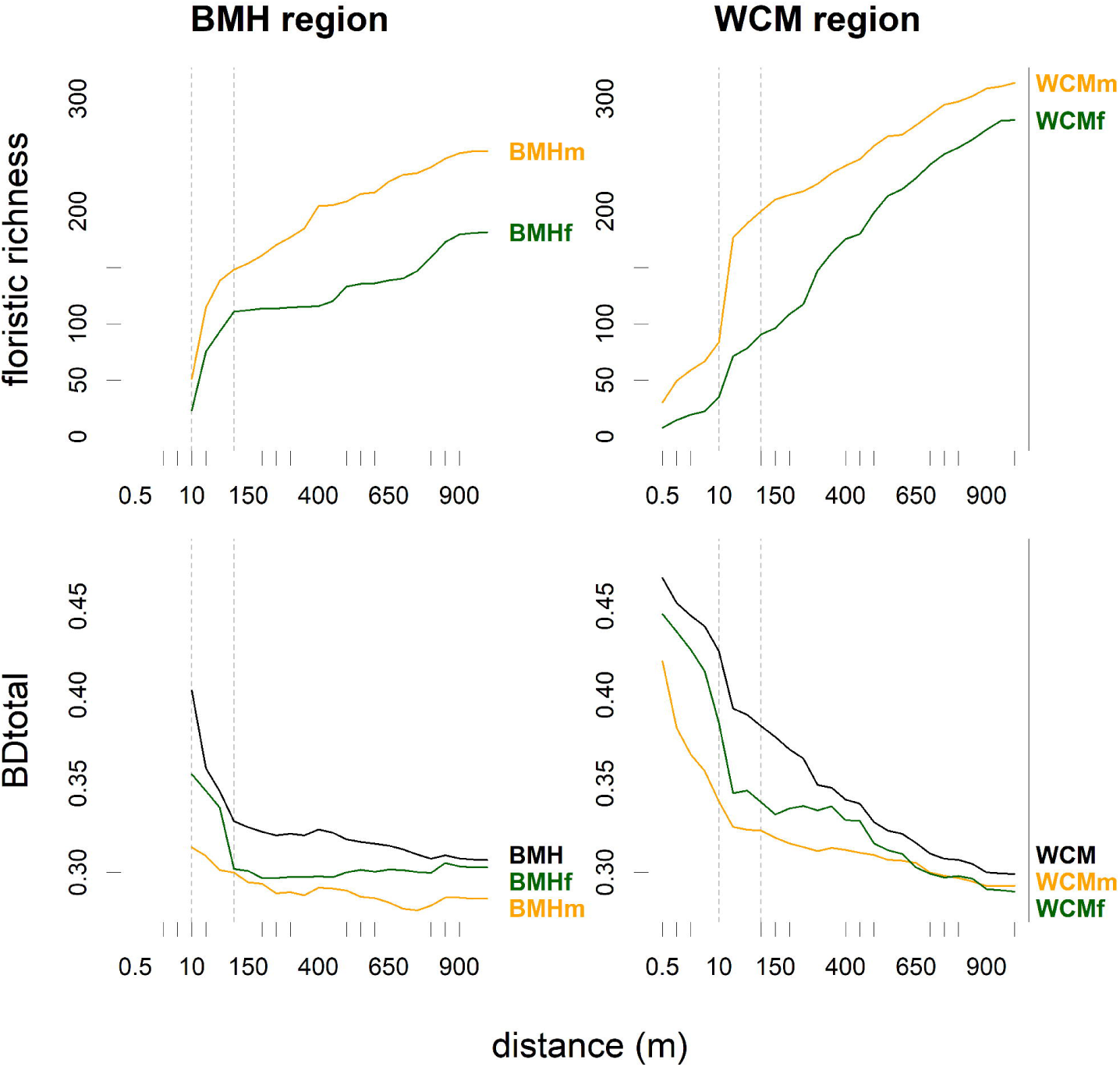
Spatial scaling of floristic alpha diversity (floristic richness) and beta diversity (BDtotal) in two study regions and their different habitats. The mean number of plant species appearing in the vegetation survey (top) and their total variance (bottom). Meadows are indicated by the yellow line, forest by the green line, and both habitats together by the black line.

BDtotal in mixed datasets of pollen and plants was always higher than in uniform datasets. Pollen and floristic BDtotals in WCM were higher than those in BMH up to a distance of 700 m. BDtotals in uniform pollen datasets varied from 0.21 in BMH meadows to 0.24 in WCM meadows (Fig. 2). Floristic BDtotal was highest in WCM forest and lowest in BMH meadows considering a distance of 40–200 m in uniform datasets. The general decreasing trend with increasing distance showed only minor exceptions, the most visible being the increase at 150 m in the WCM forest dataset (Fig. 3).

Pollen and floristic richness values were lower in forests than in meadows; however, BDtotals were higher in the forest than in meadow datasets.

### Relationship between floristic and pollen richness, calibration of alpha diversity

All datasets showed a positive correlation of pollen and floristic richness for at least some distances (Fig. 4a). Both mixed datasets and the dataset of WCM meadows showed highly significant correlations, while BMH meadows and BMH forest showed a less significant correlation, though a lower number of replications. WCM forest showed only a marginally significant correlation, despite a higher number of replications. The highest adjusted R^2^ appeared on the distances between 1.5 and 550 m, depending on the dataset (Table A3). All datasets, except WCM forest, showed two maxima of adjusted R^2^: the first one within tens of metres and the second one within hundreds of metres (450–550 m for BMH, 250–300 m for WCM). Mixed datasets had higher adjusted R^2^ than their uniform subsets, except for BMH regions between 40 and 200 m (Fig. 4a). The average distance of maximum adjusted R^2^ for six compared datasets was 286 m.

**Fig. 4:**
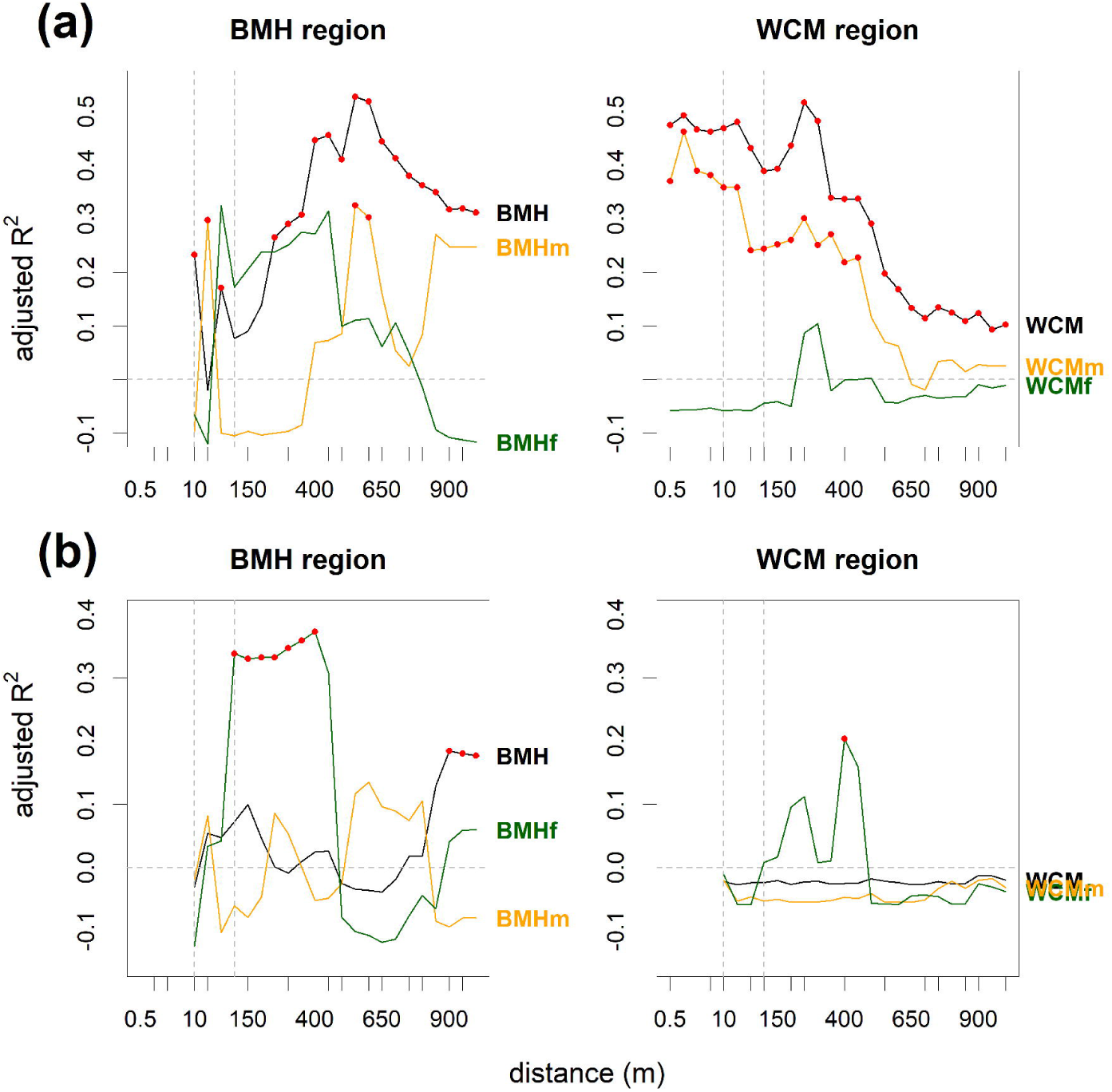
The strength of linear regression between a) pollen richness and floristic richness at different distances from sampling sites and b) local contributions of sites to pollen and floristic BDtotal at different distances from sampling sites. The black line shows the correlation for all sites, the orange line for meadow sites, and the green line for forest sites. Red dots indicate significant correlations. For more detail, see Fig. A2 and A3.

Adjusted R^2^ between pollen and floristic richness showed two general ranges of distances where the correlation was high. The first, within tens of metres from the central points, showed a correlation between pollen and plant richness values in all datasets except WCM forest (Fig. 4a). At this distance, most species naturally appear for the first time (Fig. 5). The highest correlation of pollen and plant richness values in WCM meadows is at 1.5 m within the habitat of species-rich meadows. BMH meadows had a local maximum of adjusted R^2^ at 40 m where new habitats such as forests and forest roads commonly occur. BMH forest showed the best fit at 70 m, where species confined to forest roads frequently appear (Fig. 5).

**Fig. 5:**
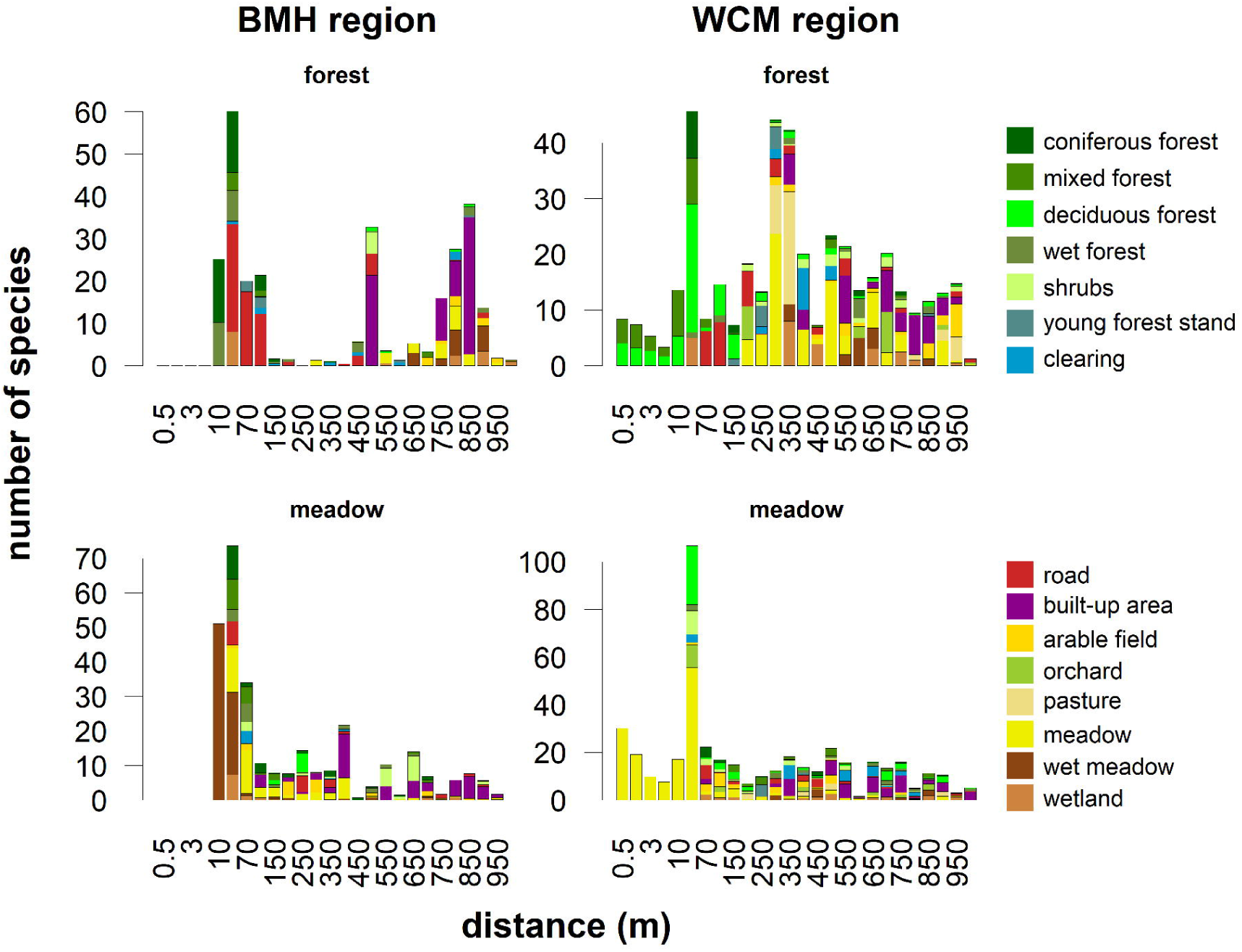
Number of new species recorded with increasing distance in different study regions and habitats, coloured according to source habitats.

The second range of maximum adjusted R^2^ values appeared between 400 and 550 in BMH meadows and between 250 and 300 m in WCM forest. Those distances match the appearance of a high number of new species in built-up areas and meadows or pastures, respectively. The local maximum of adjusted R^2^ in BMH forest at 450 m correlated with few species (< 10) from roads, clearings, and wet forests; however, a high number of species (> 30) at 500 m was accompanied by a decrease of adjusted R^2^.

Meadow datasets in both regions obtained most of the species within the first 100 m, whereas forest datasets received high numbers of species at greater distances. The floristic richness of BMH forest largely originated from human-made habitats (forest roads between 10 and 100 m, and built-up areas usually at a distance above 500 m), whereas WCM forests are enriched by meadows and other semi-natural habitats, usually at a distance above 200 m (Fig. 5).

### Relationship between pollen and floristic variance, calibration of beta diversity

The highest adjusted R^2^ between pollen and floristic BDtotal was identified at 150 m. The significant correlation appeared between 100 and 250 m, a remarkable high, but the insignificant correlation appeared between 300 and 600 m (Fig. 6a). The floristic BDtotal of WCM meadow at 150 m was lower than the floristic BDtotal of the rest of the datasets concerning the linear relationship to the pollen BDtotal (Fig. 6b). A distance of 150 m followed the steep decrease of floristic BDtotal at 10-100 m (Fig. 3), when most of the taxa appeared, and fell between both ranges of maximum adjusted R^2^ of the richness regression.

**Fig. 6:**
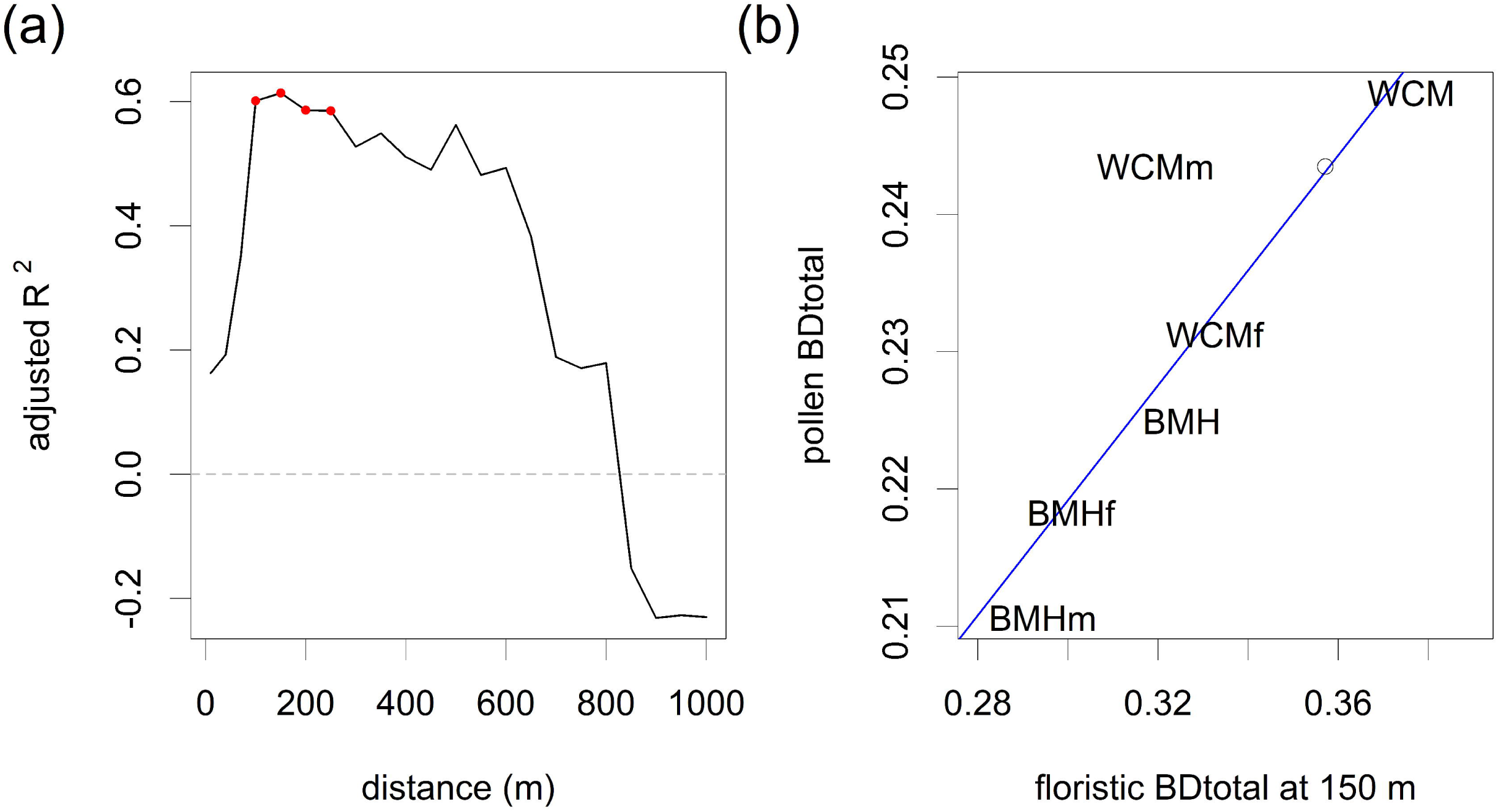
Linear regression between pollen and floristic BDtotal: a) adjusted R^2^ at different distances from sampling sites, red dots indicate significant correlations, and b) scatter plot of six datasets (text labels) for the distance of 150 m. Empty dots indicate floristic BDtotal at 6 m of WCM meadows.

Local contributions of sites to the pollen and floristic BDtotals correlated significantly in both forest datasets at 100–400 m and in the mixed dataset from the BMH region at 900–1,000 m. BMH meadows showed a positive but insignificant relationship, while meadow and mixed datasets from the WCM region did not show any relationship (Fig. 4b).

## Discussion

Our results show that the spatial distribution of plants determines the relationship between pollen and floristic richness and between pollen and floristic variance. Furthermore, the number of newly appearing species controls the correlation of richness values. Different species with respect to the whole dataset determine the correlation of variances. The appearance of new habitats at a single site often leads to an abrupt increase in species numbers. Those species may also be new for the whole dataset; in this way, the source area of pollen richness and the source area of pollen variance are linked.

### Landscape structure determines the source area of pollen richness

The source area of pollen richness, measured as adjusted R^2^ between pollen and floristic richness, is determined by numbers of new plant species appearing with increasing distance. The position of the pollen site within the landscape structure affects the order of the habitats and the sequence of appearing species. The grain size of the landscape structure is smaller in the WCM region than the BMH region; thus, the WCM region resembles a more even mosaic of habitats and the higher floristic-pollen relationship is reached at shorter distances. The same role of landscape structure was described in studies of RSAP (Bunting et al. 2004, Hellman et al. 2009a): areas with large patches of landscape have larger RSAP (Broström et al. 2005). This agreement is understandable, as both approaches seek the most appropriate distance with increasing areas of vegetation surveys. Floristic richness, assessed in our study, is a cumulative number of any new species, whereas vegetation data in RSAP approaches are cumulative abundances of main dominants, which are further distance- and dispersal-weighted. The RSAP identifies a distance at which all sites have sufficient vegetation cover of all taxa (Hellman et al. 2009b), or where all sites have all taxa present (Abraham and Kozáková 2012).

While a previous study estimated RSAP at 350–450 m (Kuneš et al. 2019), the WCM dataset allows comparison with the source area of pollen richness (250 m). Both values are similar, despite the conceptual differences mentioned above: while RSAP calculation was based on pollen/vegetation proportions of 17 taxa, here we deal with incidences of the whole spectra. Moreover, vegetation structure for the range between 10 and 1,000 m was recorded independently in the two studies. We suggest that this robustness indicates the significant effect of landscape structure on the pollen–vegetation relationship.

A critical characteristic of the landscape mosaic, affecting the source area of pollen richness, appears to be the number of species per patch rather than its area (Fig. A4). The appearance of open habitats within forest led to the increase of species numbers and the approaching of local maxima of adjusted R^2^ in both regions. While in BMH forest the appearance of forest roads at about 70 m was crucial, meadows and orchards at about 250 m played a similar role in WCM forest. Other studies in semi-open landscapes which found a high correlation between pollen richness and landscape openness (Meltsov et al. 2013, Matthias et al. 2015) corroborate this idea. Source areas of pollen richness in forests of both regions fell close to the transition from species-poor to species-rich habitats. Meadow datasets, i.e., cases where pollen sites were in the species-rich habitats, provided additional insight regarding the compositional similarity between central habitats and those in the surrounding landscape. In BMH meadows, there was an increased correlation of floristic and pollen data at 400 and 550 m. This distance relates to the increase in the number of species due to the frequent transition of meadow complexes to shrubby habitats and built-up areas. We generalise the “landscape openness” finding of Matthias et al. (2015) and Meltsov et al. (2013) to the more general “species-rich patches”.

### Effect of biodiversity hotspot on source area of pollen diversity

Most of the species in WCM meadows appeared within the first 40 m in the dominant habitat of extremely species-rich steppic grasslands (Fig. 5). Even though habitats such as built-up areas and roads appearing beyond 40 m can be potentially species-rich and compositionally different to those grasslands, the number of new species appearing between 100 and 1,000 m is small (usually below 20). It is apparent that high fine-scale floristic diversity suppresses the influence of the surrounding landscape on pollen richness and decreases the source area of pollen richness. The relationship is stronger at 1.5 m (0.46) than at 250 m (0.3).

The strong effect of high pollen richness in WCM meadows is also visible in the comparison of pollen and floristic variance. At 150 m, WCM meadows had much lower floristic variance than the other datasets. Floristic variance in WCM meadows corresponding to the pollen variance and the pattern of the other datasets lay at 6 m (Fig. 6b). Again, this may be caused by the high fine-scale diversity of the meadows, which include most pollen types present in the surrounding landscape. Only a few new species appeared in broader surroundings. WCM meadow sites are too similar at 150 m than other analysed habitats. Indeed, a similar result, showing compensation of extremely high alpha diversity by low beta diversity, has already been reported from the White Carpathians (Michalcová et al. 2014).

### Floristic reference of pollen-based beta diversity

Estimating the source area of pollen variance as a regression of pollen and floristic variance implies that the resulting distance of 100-250 m represents all datasets. Though they differ in species richness, openness, and habitats of origin, the relationship between variances is fairly linear. The only exception is the biodiversity hotspot of WCM meadows mentioned above. It shows that the spatial scale at which the pollen variance corresponds to the floristic variance is dataset-specific. The linearity and the significance of the variance relationship within the rest of the datasets indicate certain robustness and possible applicability to a variety of fossil records from peat bogs.

The mechanism of establishing the source area of pollen variance was similar to that mentioned for the source area of pollen richness. The appearance of new habitats with new species (Fig. 5), like open habitat for forest sites (WCM forest) or built-up areas for meadow sites (BMH meadow), caused small to negligible increases of floristic variance. Moreover, the high, yet insignificant relationship of the variances at the distance 250-600 m (Fig. 6a) corresponds to the distance of the second range of fit between richness (Fig. 4a).

The link between pollen and total floristic variance is underlined by the relationship between the amount of variance contributed by individual sites and the total variance. Indeed, these amounts in pollen and floristic data are significantly correlated. Distances of the high correlation of local contribution to total variance are related to the source area of pollen richness of individual datasets. In WCM forest and BMH forest, the increase of correlation of local contribution to beta diversity usually follows (or precedes) the richness correlation by a single ring (Fig. 4).

Beta diversity understood as directional turnover (temporal or spatial) belongs to more frequent and widespread measurements in pollen analysis (Figueroa-Rangel et al. 2010, Connor et al. 2019) than beta diversity as a non-directional variation. Indeed, the temporal dynamics is the main objective of pollen analysis. According to Nieto-Lugilde et al. (2015) pollen-based turnover correlates with forest-inventory-based turnover. We extend this finding from woody taxa to all species and from directional turnover to non-directional variance. Moreover, forest sites with high contributions to pollen beta diversity also show a high contribution to floristic beta diversity (Fig. 4b).

### Strengths and limitations

We applied the presence-absence transformation of abundances of pollen data instead of widely used pollen proportions (Birks et al. 2016). This transformation may increase the effect of very rare taxa, sometimes appearing in a single grain due to long-distance transport. While we admit that our datasets contain such pollen components (*Ambrosia artemisiifolia*-type, *Ostrya, Castanea*), their presence was perhaps balanced by rare ornamental plants recorded in the plant survey, which were missing in the pollen record. In this light, the significance of the relationship between pollen-based beta diversity and floristic data is a remarkable result of this study and provides a rigorous reference for pollen-based macroecological studies (e.g. Šizling et al. 2016).

Disentangling the causal connection in any pollen–vegetation relationship requires the use of pollen–plant translation tables (Birks et al. 2016) and the comparison of pollen and vegetation patterns by individual taxa. Here, we report metrics inferred from all-species datasets. Without relying on any particular pollen dispersal model, we assumed that the correlation of pollen- and floristic-based metrics is caused by pollen transport. Studies reconstructing vegetation from pollen usually assume that the over-canopy component is the major component of pollen transport and do not consider the local herb layer in the forest (Prentice 1985). However, we identified a positive correlation between pollen and floristic richness at 70 m in BMH forest, matching the distance at which many open-habitat species, confined to forest roads and other openings, first appeared. If the plant individuals recorded in these habitats are direct sources of the recorded pollen, then pollen transported through the forest interior also has to be considered (Tauber 1967). Our data also indicated, however, that forest serves as a barrier against pollen transport. While many new plant species recorded in the built-up areas in the BMH region caused a decrease of adjusted R^2^ at 500 m in forest habitats, the opposite pattern was recorded in meadow sites in the same region. Open sites of WCM meadows also show a relationship between dispersal abilities of pollen and the source area of pollen richness. The distance of 1.5 m in the herb plants-dominated WCM meadows corresponds to the taxon-specific source area of pollen for herbs. In this regard, insect-pollinated strategy with low pollen productivity makes source areas of individual plants very small (Shaw and Whyte 2020).

We had to consider the trade-off between data quality and the feasibility of the vegetation survey. For the range of radii 10–100 m, we mapped vegetation in the field with the help of aerial photographs; however, radial relevés for each ring might have been better (Shaw and Whyte 2020). In the case of homogeneous vegetation, our method produced a single or a few large polygons, and thus most species appeared “in waves” along the transects (from 10 to 40 m in WCM meadows). Heterogeneous vegetation resulted in higher numbers of polygons and species appeared gradually along the transect there (BMH forest, Fig. 3).

Our survey methodology concentrates sampling effort around central points, which then decreases with increasing radius, though not smoothly. Change from polygon to transect method at 100 m is noticeable in the trend of floristic richness in the BMH region, which slightly slowed down beyond 100 m. Our survey methodology, emphasising a ‘pollen perspective’ of vegetation, may be contrasted with independent data sources, to illustrate the completeness of our data. The most complete database of Czech flora PLADIAS includes information on 1,477 species in 15 mapping squares covered by our survey in the BMH region and 2,045 species in 14 squares in the WCM region (Wild et al. 2019). This means that in both regions, we recorded close to 54% of the known regional species pool. This is quite good result, considering the incomplete spatial coverage of our survey. Moreover, the close agreement between the two regions speaks for consistency in data quality between the datasets.

## Conclusions

There was a consistent positive relationship between pollen and floristic richness in both open and forest habitats in the two study regions. The strongest correlations mostly appeared at two spatial scales: within tens and hundreds of metres. These distances are controlled by numbers of newly appearing species along the transects, linked to landscape structure and the mean of the pollen transport. However, causality between the appearance of species and pollen taxa at individual sites remains tangled.

Our results support the hypothesis that a higher variance of datasets covering both open and forest habitats may contribute to a higher correlation within the forest and open-habitat subsets. Regarding the application value of these results for the interpretation of fossil records, we suggest that pollen richness reconstructions from sites with different openness or richness should be compared, since the non-significant relationships recovered in both forest datasets indicates some limitations of the method.

Total variation was already used as a measure of spatial beta diversity in paleoecology (Winegardner et al. 2017). We encourage its broader use in palynology and macroecology. Our field data prove that pollen-based beta diversity has a significant relationship to floristic beta diversity. The source area of pollen variance appears to be dataset-specific. Our results, pointing close to 150 m, represent the mean spatial reference for pollen-based beta diversity inferred from moss polsters at open, forest, species-poor, and species-rich sites. This result is also underlined by a significant correlation between pollen and floristic indices of local contributions of sites to the variation in both forest datasets. Decomposition of beta diversity on the level of individual sites or species is one of the advantages of the total variance (Legendre and De Cáceres 2013) and it has already been shown that it may provide additional insights to the fossil datasets (Connor et al. 2019).

## Declarations

### Funding

This study was financed by the Czech Science Foundation (Grant No. 16-10100S). Authors affiliated with the Institute of Botany were further supported by the long-term developmental project of the Czech Academy of Sciences (RVO 67985939).

### Author contributions

PK, JR, VA conceived initial idea; JR, OV, VA developed methodology of vegetation data sampling; VA analysed of data and drafted the manuscript; all authors collected the data and commented the manuscript.

### Conflicts of interest

The authors declare no competing interests.

### Permit(s)

Permission to enter the nature reserves was given by the Nature Conservation Agency of the Czech Republic.

### Data availibility

pollen data will be available in the Neotoma Palaeoecological database Code availibility – code to reproduce the numerical analysis will be available at https://github.com/vojtechabraham/SpatialScalingPollenDiversity/

## Acknowledgements

The Nature Conservation Agency of the Czech Republic is acknowledged for granting permission to access the nature reserves. The authors are grateful to the following colleagues who kindly helped during any stage of the fieldwork: Přemysl Bobek, Zita červenková, Pavel Daněk, Pavel Dřevojan, Michelle Farrell, Radim Hédl, Markéta Chudomelová, Kryštof Chytrý, Radka Kozáková, Pavel Novák, and Helena Prokešová.

## Notes

https://github.com/vojtechabraham/SpatialScalingPollenDiversity

### Competing Interest Statement

The authors have declared no competing interest.

