## Supplementary Material for "Spatial scaling of pollen-based alpha and beta diversity within forest and open landscapes of Central Europe"

### Appendix 1

We are aware of the problem of different detection probability when some samples are dominated by high pollen producers. Treatment of pollen data, in this case, proposed by Odgaard (1999), is the division of pollen counts by pollen productivity. We picked mean values for the northern Hemisphere (Table A1; Wieczorek and Herzschuh 2020), which contain values calculated by us from the same pollen data in the WCM region (Kuneš et al. 2019). After adjusting pollen counts by PPEs, we decreased the raw and PPE-adjusted dataset to the sum of 350 grains.

BMH forest and BMH meadow do not show any difference between raw and adjusted pollen counts. Datasets with all sites clearly show a worse fit after PPE adjustment. The WCM forest slightly improves and WCM meadows are the only dataset that clearly improves after PPE adjustment (Fig. A1).

These results led us to the decision to work through the whole paper with only raw pollen counts. This confirms the finding of Felde et al. (2016), that the problem of different detection probability matters only in the boreal zone, where high pollen producers appear and where adjustment of pollen counts by PPEs led to the improvement of the relationship between pollen and floristic richness (Odgaard 1999, Felde et al. 2016, Reitalu et al. 2019).

Table A1: Pollen productivity estimates (Wieczorek and Herzschuh 2020)

| Taxon | PPE |
| --- | --- |
| *Abies* | 6.88 |
| *Alnus* | 7.46 |
| *Betula pubescens* type | 4.4 |
| *Carpinus betulus* | 4.52 |
| *Corylus* | 1.97 |
| *Fagus* | 1.96 |
| *Fraxinus excelsior*-Typ | 1.25 |
| *Juniperus*-Typ | 9.8 |
| *Picea* | 2.29 |
| *Pinus sylvestris*-Typ | 10.47 |
| *Quercus* | 3.33 |
| *Tilia* | 1.17 |
| *Ulmus* | 7.32 |

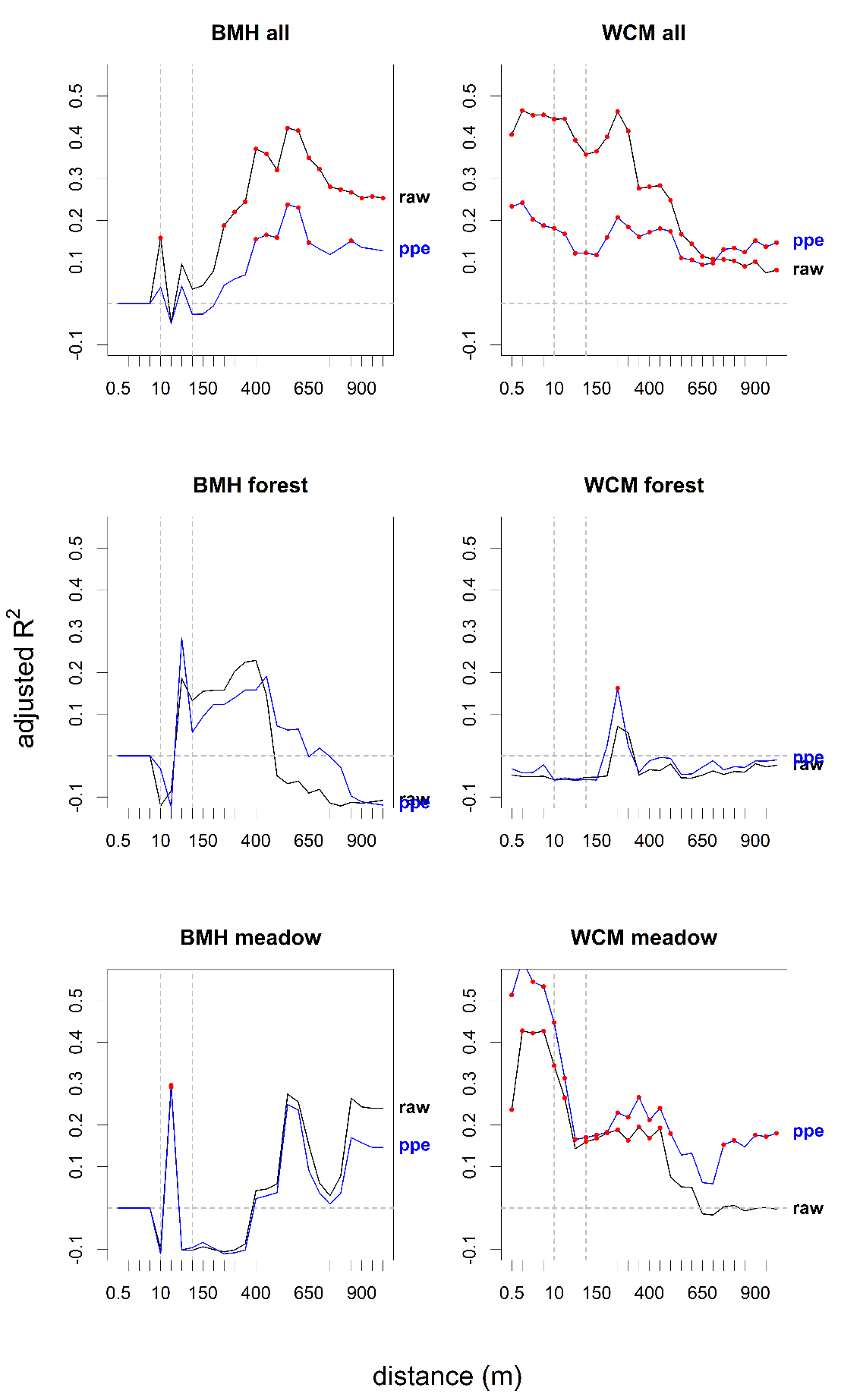

Fig. A1: The strength of linear regression between pollen richness and floristic richness at different distances from sampling sites. The black line shows a correlation for raw pollen counts; the blue line shows pollen counts adjusted by PPE. Red dots indicate significant correlations.

Table A2: Sites.

| **reg** | **id** | **name** | **type** | **sampl. date** | **longitude** | **latitude** | **altitude** | **pollen sample material** |
| --- | --- | --- | --- | --- | --- | --- | --- | --- |
| BMH | 3 | Plíčky | open | 29-MAR-2016 | 15.97328 | 49.56648 | 600 m | *Calliergonella cuspidata* |
| BMH | 4 | Louky u Černého lesa | open | 29-MAR-2016 | 15.94290 | 49.58620 | 581 m | mosses |
| BMH | 6 | Račín | forest | 29-MAR-2016 | 15.87842 | 49.61474 | 665 m | *Sphagnum* sect. *Squarrosum* |
| BMH | 7 | Vepřová-Žlábek | forest | 29-MAR-2016 | 15.83664 | 49.62626 | 656 m | *Sphagnum* cf. *Girgensohnii* |
| BMH | 10 | Suché Kopce | open | 29-MAR-2016 | 15.89515 | 49.68509 | 661 m | *Sphagnum* sect. *palustria* |
| BMH | 11 | Pihoviny | open | 30-MAR-2016 | 15.97139 | 49.65886 | 676 m | *Sphagnum teres* |
| BMH | 12 | Kocanda | open | 30-MAR-2016 | 15.98694 | 49.68227 | 644 m | Sphagnum sect. *palustria* |
| BMH | 13 | Porostliny | open | 30-MAR-2016 | 16.06020 | 49.76057 | 590 m | *Sphagnum* cf. *Angustifolium* |
| BMH | 14 | Bahna | open | 30-MAR-2016 | 15.99222 | 49.75349 | 655 m | *Sphagnum* sect. *palustria* |
| BMH | 15 | Ratajské rybníky | open | 30-MAR-2016 | 15.93400 | 49.76956 | 592 m | *Sphagnum* sect. cf. *Cuspidata* |
| BMH | 16 | Zubří | open | 30-MAR-2016 | 15.79089 | 49.77894 | 624 m | *Sphagnum* cf. *teres* |
| BMH | 17 | Nový Rybník | open | 30-MAR-2016 | 15.81989 | 49.80366 | 572 m | *Sphagnum teres* + *S*. Sect. *Subsecunda* |
| BMH | 18 | Stropnická cesta | forest | 30-MAR-2016 | 16.11238 | 49.74904 | 727 m | *Sphagnum* cf. *Girgensohnii* |
| BMH | 19 | Žižkov | forest | 30-MAR-2016 | 16.13217 | 49.73117 | 760 m | *Sphagnum* cf. *Girgensohnii* |
| BMH | 20 | Samotín | open | 30-MAR-2016 | 16.06899 | 49.65377 | 706 m | *Sphagnum* cf. *Fallax* |
| BMH | 21 | Chlum | forest | 31-MAR-2016 | 15.85758 | 49.73012 | 569 m | *Sphagnum* cf. *Girgensohnii* |
| BMH | 22 | Míšek | forest | 31-MAR-2016 | 16.03064 | 49.74724 | 683 m | *Sphagnum* cf. *Girgensohnii* |
| BMH | 23 | Knížecí studánka | forest | 31-MAR-2016 | 16.07440 | 49.71121 | 724 m | *Sphagnum* sect. *palustria* |
| BMH | 24 | Pod Šindelným vrchem | forest | 31-MAR-2016 | 15.95803 | 49.67219 | 739 m | *Sphagnum* cf. *Girgensohnii* |
| BMH | 25 | Rampoltův mlýn | forest | 31-MAR-2016 | 16.01390 | 49.66028 | 704 m | *Sphagnum* cf. *Riparium* |
| BMH | 26 | Brožova skála | forest |  | 16.01675 | 49.62796 |  | mosses |
| WCM | 2 | B1 | open | 19-MAY-2013 | 17.53333 | 48.88450 |  | mosses |
| WCM | 4 | B10 | open | 27-JUN-2013 | 17.52458 | 48.83287 |  | mosses |
| WCM | 6 | B11 | open | 09-JUL-2013 | 17.55803 | 48.87033 |  | mosses |
| WCM | 8 | B12 | open | 10-JUL-2013 | 17.43011 | 48.86400 |  | mosses |
| WCM | 10 | B13 | open | 11-JUL-2013 | 17.42975 | 48.84800 |  | mosses |
| WCM | 12 | B14 | open | 12-JUL-2013 | 17.40272 | 48.85531 |  | mosses |
| WCM | 14 | B15 | open | 19-JUL-2013 | 17.44619 | 48.84497 |  | mosses |
| WCM | 36 | B16 | open | 21-MAY-2014 | 17.28043 | 48.85752 |  | mosses |
| WCM | 38 | B17 | open | 19-JUN-2013 | 17.48550 | 48.90047 |  | mosses |
| WCM | 39 | B18 | open | 05-JUN-2014 | 17.39347 | 48.87321 |  | mosses |
| WCM | 40 | B19 | open | 04-JUN-2013 | 17.42745 | 48.87539 |  | mosses |
| WCM | 20 | B2 | open | 20-MAY-2013 | 17.59433 | 48.90500 |  | mosses |
| WCM | 41 | B20 | open | 03-JUN-2013 | 17.45899 | 48.88813 |  | mosses |
| WCM | 23 | B3 | open | 21-MAY-2013 | 17.60039 | 48.88214 |  | mosses |
| WCM | 25 | B4 | open | 21-MAY-2013 | 17.61483 | 48.89183 |  | mosses |
| WCM | 27 | B5 | open | 22-MAY-2013 | 17.65044 | 48.89639 |  | mosses |
| WCM | 29 | B6 | open | 23-MAY-2013 | 17.67744 | 48.91792 |  | mosses |
| WCM | 31 | B7 | open | 23-MAY-2013 | 17.72650 | 48.88992 |  | mosses |
| WCM | 33 | B8 | open | 23-MAY-2013 | 17.63767 | 48.92156 |  | mosses |
| WCM | 35 | B9 | open | 26-JUN-2013 | 17.32711 | 48.86350 |  | mosses |
| WCM | 17 | L1 | forest | 18-JUN-2012 | 17.60442 | 48.92819 |  | mosses |
| WCM | 24 | L10 | forest | 04-AUG-2012 | 17.44664 | 48.85539 |  | mosses |
| WCM | 32 | L11 | forest | 05-AUG-2012 | 17.34517 | 48.84572 |  | mosses |
| WCM | 34 | L12 | forest | 06-AUG-2012 | 17.31531 | 48.84117 |  | mosses |
| WCM | 3 | L13 | forest | 06-AUG-2012 | 17.32297 | 48.80853 |  | mosses |
| WCM | 5 | L14 | forest | 06-AUG-2012 | 17.52731 | 48.81708 |  | mosses |
| WCM | 11 | L16 | forest | 07-AUG-2012 | 17.48517 | 48.82239 |  | mosses |
| WCM | 16 | L17 | forest | 07-AUG-2012 | 17.39308 | 48.80753 |  | mosses |
| WCM | 13 | L18 | forest | 29-JUL-2013 | 17.44197 | 48.83097 |  | mosses |
| WCM | 15 | L19 | forest | 06-AUG-2013 | 17.39228 | 48.79989 |  | mosses |
| WCM | 18 | L2 | forest | 20-JUN-2012 | 17.64928 | 48.91847 |  | mosses |
| WCM | 21 | L20 | forest | 20-MAY-2014 | 17.60472 | 48.92004 |  | mosses |
| WCM | 1 | L3 | forest | 21-JUN-2012 | 17.49997 | 48.83731 |  | mosses |
| WCM | 28 | L4 | forest | 21-JUN-2012 | 17.54692 | 48.87761 |  | mosses |
| WCM | 30 | L5 | forest | 23-JUL-2012 | 17.56997 | 48.89289 |  | mosses |
| WCM | 9 | L6 | forest | 24-JUL-2012 | 17.50275 | 48.82275 |  | mosses |
| WCM | 26 | L7 | forest | 25-JUL-2012 | 17.59197 | 48.89292 |  | mosses |
| WCM | 19 | L8 | forest | 03-AUG-2012 | 17.54653 | 48.89542 |  | mosses |
| WCM | 22 | L9 | forest | 04-AUG-2012 | 17.44522 | 48.87050 |  | mosses |

Table A3: Relationship of floristic and pollen richness within the datasets.

| **region** | **dataset** | **n sites** | **distance max fit (m)** | **adjusted R^2^** | **p_value** |
| --- | --- | --- | --- | --- | --- |
| BMH | forest | 10 | 70 | 0.3235547 | 0.050 |
| BMH | open | 11 | 550 | 0.3201134 | 0.041* |
| BMH | all | 21 | 550 | 0.525054 | 0.000*** |
| WCM | forest | 19 | 300 | 0.1037916 | 0.097 |
| WCM | open | 20 | 1.5 | 0.4623916 | 0.001*** |
| WCM | All | 39 | 250 | 0.5167158 | 0.000*** |

significance levels: ***<0.001, *<0.05

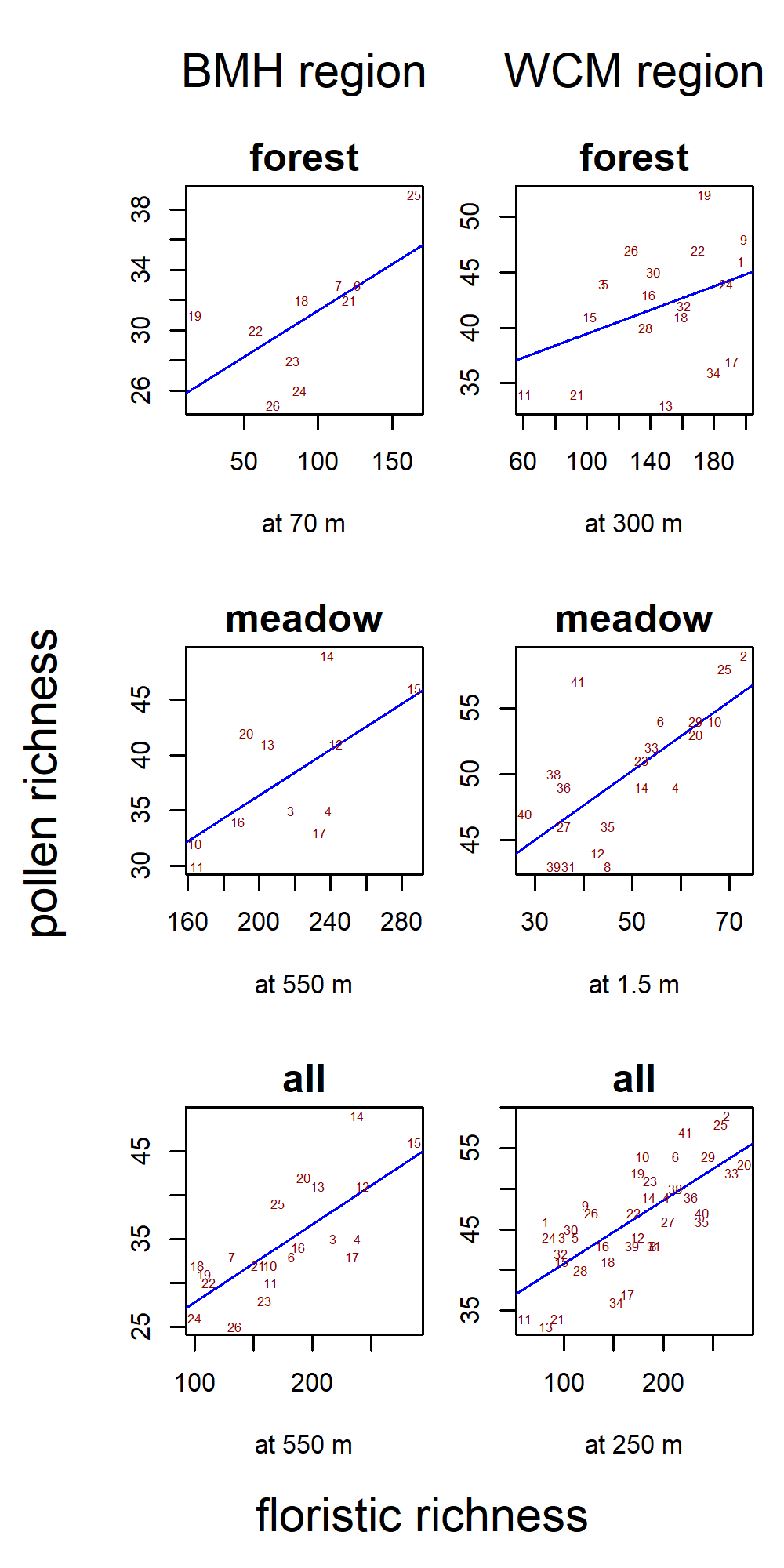

Fig. A2: The relationship between pollen richness and floristic richness at the distances with maximum overall correlation (see Table A3). Red numbers are sites’ IDs (see Table A2).

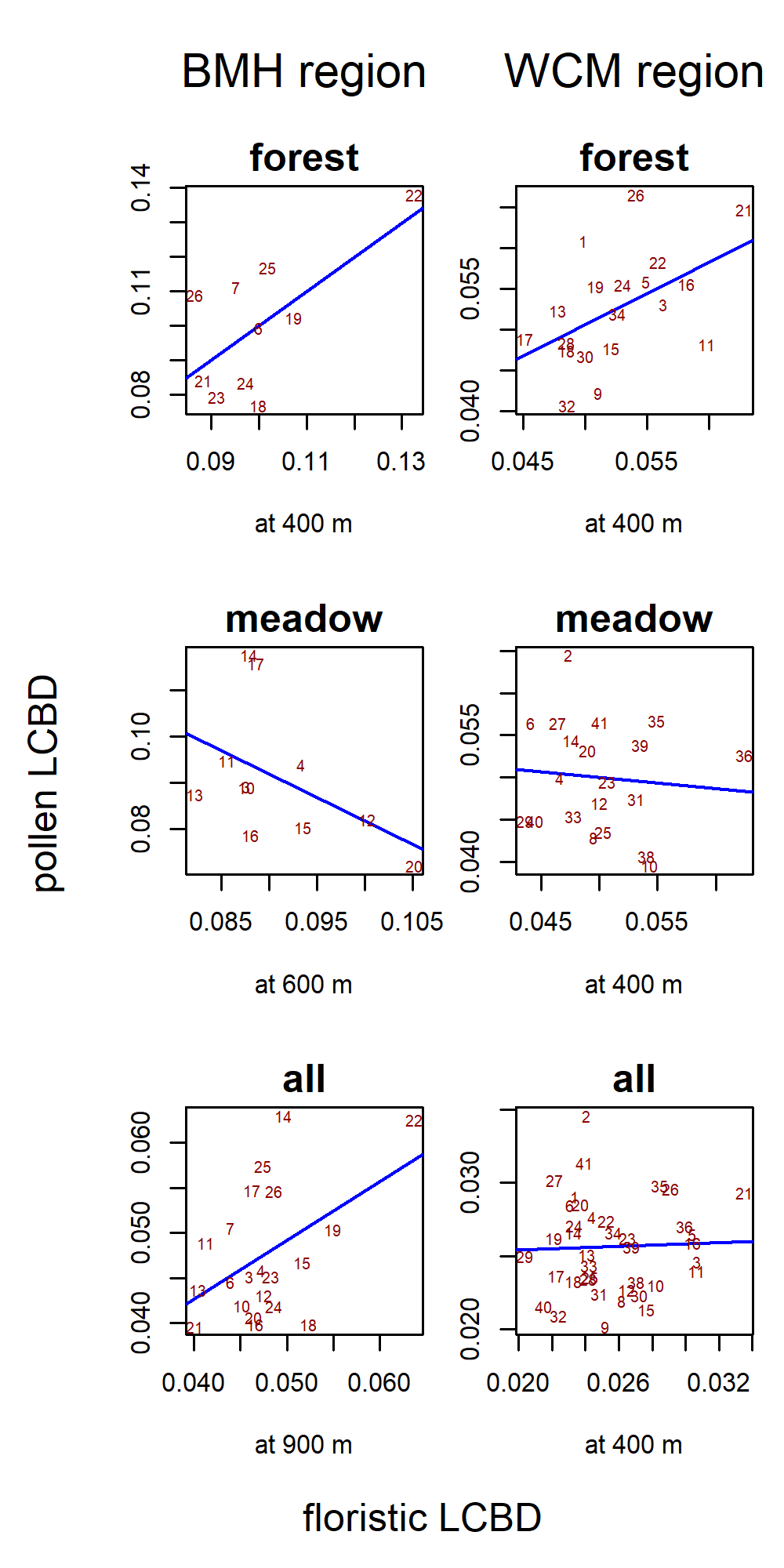

Fig. A3: The relationship between the local contribution of sites to pollen beta diversity and the local contribution of sites to floristic beta diversity at distances with maximum overall correlation (see Fig. 4b). Red numbers are sites’ IDs (see Table A2).

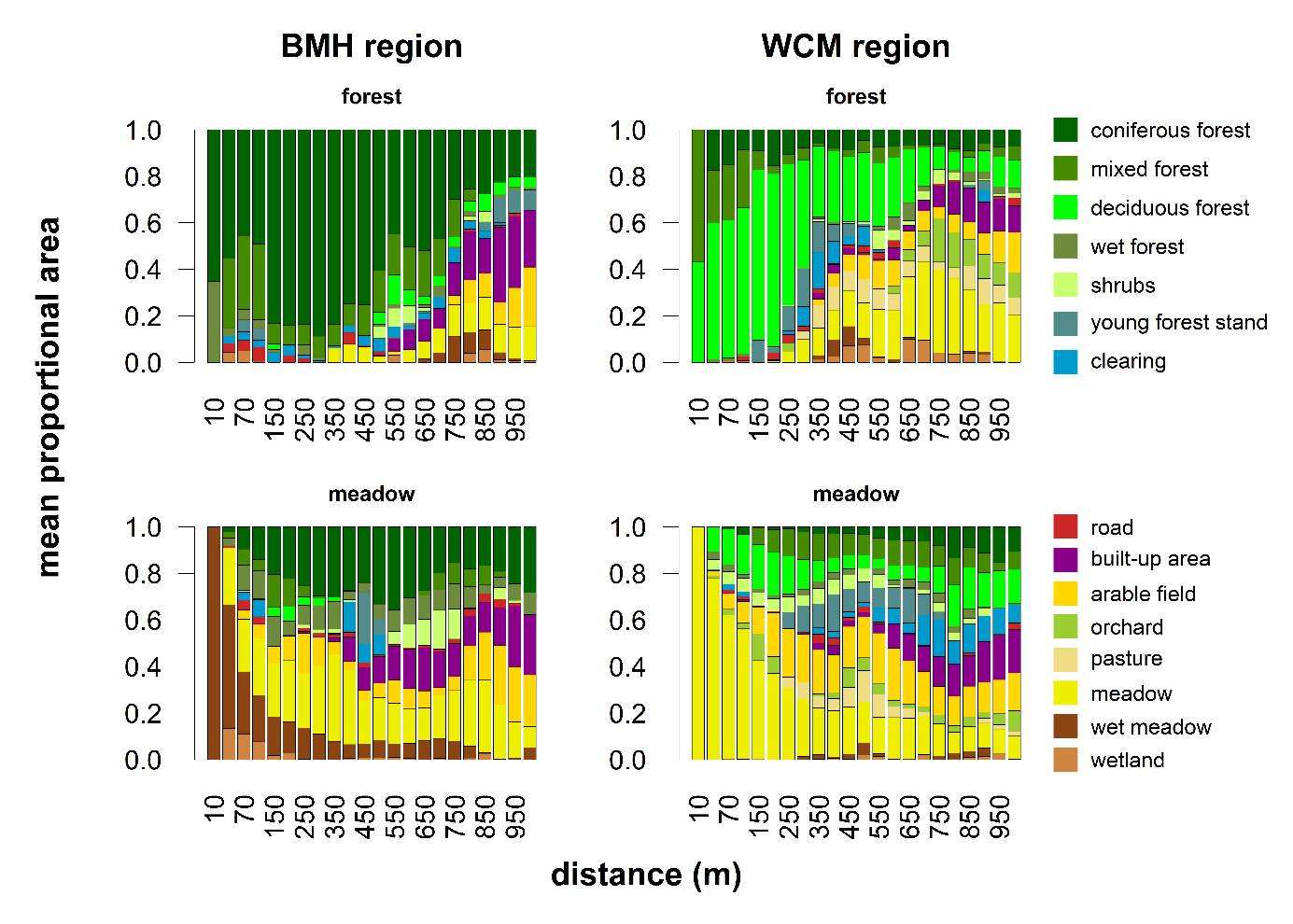

Fig. A4: Cumulative composition of habitats in the rings of the uniform datasets. The proportion of habitats corresponds to the relative area of each polygon within the ring.
